## Supplementary Information for "One-pot Golden Gate Assembly of an avian infectious bronchitis virus reverse-genetics system"

Supplementary Table 1. Domesticating Mutations.

| Location | Mutation | Codon | Gene | Fragment |
|---|---|---|---|---|
| 699 | T → A | GGT (Gly) → GGA (Gly) | nsp2 | D388-F2 |
| 855 | T → A | TCT (Ser) → TCA (Ser) | nsp2 | D388-F2 |
| Beau-CK 20,907 | C → T | ACC (Thr) → ACT (Thr) | Beau-CK spike | D388-F10-Beau(S) |

Silent mutations were made to remove native two BsaI sites from the IBV genome and Beau-CK spike gene sequence.

Supplementary Figure 1.

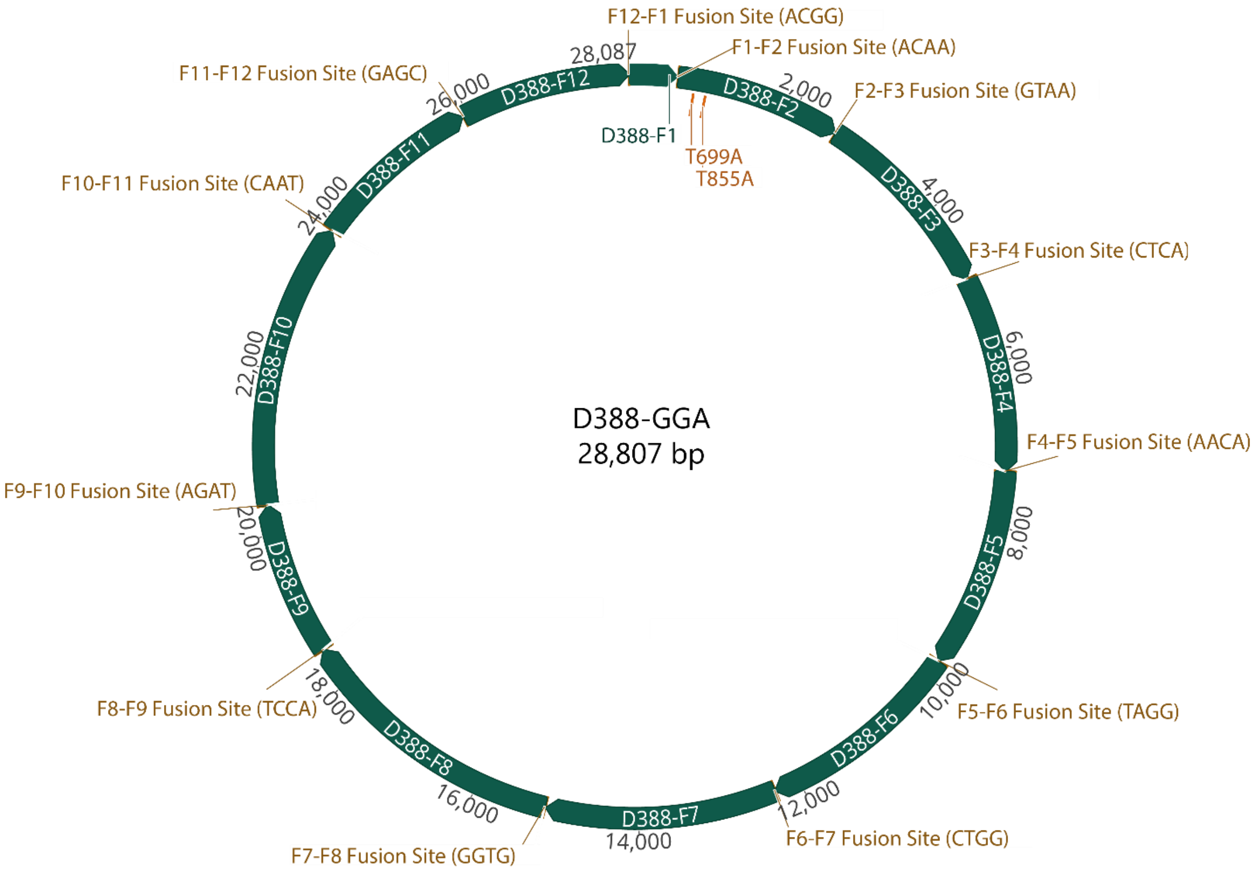


Schematic of the IBV D388-GGA construct noting the location of fusion sites and silent mutations used to remove native BsaI recognition sites. A Genbank file of the full genome with the annotations is also provided as part of the Supplementary Information.

Supplementary Table 2. Sequences of assembly fragments. All fragments were held as plasmids in a pUC57-mini Bsa-Free backbone from Genscript. For each fragment, lowercase font indicates Type IIS sites and spacers and UPPERCASE indicates the insert sequence.

| **Fragment** | **Inserted Sequence** |
|---|---|
| D388-F1 | ggctacggtctctACGGCTACGCGGCCGCCTGCAGGGCCCGGCGCGCCGGATCCATTAATACGACTCACTATAGGGACTTAAGTGTGATATAAATATATATCATACATACTAGCCTTGCGCTAGATTTCCAACTTAACAAAACGGACTTAAATACCTACAGCTGGTCCCTATAGGTGTTCCATTGCAGTGCACTTTAGTGCCCTGGATGGCACCTGGCCACCTGTCAGGTTTTTGTTATTAAAATAACATTGTTGCTGGTATCACTGCTTGTTTTGCCGTGTCTCACTTTATACATCCGTTGCTTGGGCTACCTAGTATCCAGCGTCCTACTGGCGTTGTGGTCGGTCTGAGTGCGAAGAACCTCTGGTTCATCTAGCGGTACGCGGGTGTGTGGAAGTAGCGCTTCAGACGTACCGGTTCTGTTGCGTGAAATACGGGGTCACCTCCCCCCACATACCTCTAAGGGCTTTTGAGCCTAGCGTTGGGCTACGTTCTCGCACAAGGTCGGTTATACGACGTTTGTAGGGGGTAGTGCCAAACAACCCCTGAGGTGACAGGTTCTGGTGGTGTTTAGTGAGCAGACATACAATAGACAGTGACAAggagaccgtagcc |
| D388-F2 | ggctacggtctccACAACATGGCTTCAAGCCTAAAACAGGGAGTATCTCCCAAACCAAGGAATGTCATTCTTGTTGCCAAAGACATTCCCGAACAACTCTGTGACGCTTTGTTCTTCTATACGTCACATAACCCTAAGGATTACGCTGAAGCCTTTGCCCTTAGGCAGAAGTTTGACCGTAATCTGCAGACTGGGAGGCAGTTTAAATTTGAAACTGTGTGTGGACTCTTCCTCTTGAAAGGGGTTGACAAAATAACACCTGGTGTCCCAGCAAAAGTTCTAAAAGCCACTTCTAAGTTGGCAGATTTAGAAGACATCTTTGGTGTCTCTCCCTTTGCACGGAAGTACCGTGATCTGTTGAAAACAGCATGCCAGTGGTCACTTACAGTAGACACACTGGATGCTCGTGCACAAACTCTTGATGAAATTTTTGACCCCACCGAAATACTTTGGCTTCAGGTGGCTGCAAAAATTCAAGTTTCAGCTATGGCAATGCGCAGGCTCGTTGGAGAAGTAACTGCAAAAGTCATGGATGCTTTGGGCTCAAATTTGAGTGCTCTCTTTCAAATTGTTAAAGAACAAATAGCCAGAATTTTTCAAAAAGCCCTGGCTATTTTTGAGAGTGTGAATGATTTACCACAGCGTATTGCAGCACTAAAAATGGCTTTTGCCAAGTGTGCGAAATCAATTACAGTTGTGGTTGTTGAAAAGACTCTTCTTGTTAAAGAGTTTGCTGGAACTTGTCTTGCAAGCATTAATGGTGCTATTGCAAAGGTTTTTGAAGAACTGCCAACTGGCTTTATGGGTGCCAAAATTTTCACAACACTTGCCTTCTTTAAGGAAGCAGCTGTGAAAATTGTGGAAAATATACCAAATGCACCAAGAGGTACTAGAGGTTTTGAAGTCGTTGGTAACGCCAAGGGAACGCAAGTTGTTGTGCGTGGCATGCGAAATGATTTAACTCTGCTCGACCAAAAAGCTGACATTCCTGTTGAGAAAGAAGGTTGGTCTGCAATTCTTGAAGGACATCTGTGTTATGTCTTTAGGAGTGGGGACCGGTTTTATGCGGCACCTCTTTCTGGAAATTTTGCATTGCATGATGTACATTGCTGCGAGCGTGTTGTCTGTCTTTCTGATGGTGTAACACCAGAGATAAATGATGGACTCATTCTTGCTGCAATTTACTCATCTTTTAGTATATCAGAACTTGTGGCAGCTCTTAAAAAGGGTGAACCATTCAAGTTCTTGGGCCATAAATTTGTGTACGCGAAGGATGCAGCAGTGTCGTTTACTTTAGCGAAGGCTGCCACTATTGCAGATGTCCTAAAGCTGTTTCAATCAGCGCGTGTGAAGGCAGAAGATGTTTGGTCTTCATTTACTGAAAAGTCTTTTGAGTTCTGGAAGATCGCATATGGAAAAGTGCGCAATCTTGAAGAATTTGTTAAGACTCATTTTTGTAAGGCTCAAATGTCAATTGTTGTTCTAGCAGCAGTGCTTGGAGAAGGCATTTGGCATCTTGTCTCTCAGGTCATCTATAAATTAGGTGGTCTTTTCACTAAAGTTGTTGATTTTTGTGAGAAACATTGGAAAGGGTTTTGTGCACAGTTGAAGAAGGCTAAGCTCATTGTGACTGAAAATCTTTGTGTTTTAAAAGGAGTTGCACAGCACTGTTTCCAACTACTGCTGGATGCGATATACTCTATGTACAAGAGCTTTAAAAAGTGTGCACTTGGTAGAGTCCATGGAGATCTTCTCTTCTGGAAAGGGGGTGTGCATAAAATTGTTCAAGATGGCGATGAAGTTTGGTTTGACGCCATTGATAGTGTTGATGTTGAAGATCTGGGTGTTGTCAAAGAAAAGCCTATAGATTTTGAGGTTTGTGAAGACATAACACTTCCAGAAAATCAACCTGGTCATATGGTTCAAATCGAGGATGACGGAAAGAACTATATGTTCTTCCGCTTCAAAAGGGATGAGAACATCTACTATACACCAATGTCTCAACTTGGTGCAATTAATGTAGTTTGCAAAGCAGGCGGTAAggagaccgtagcc |
| D388-F3 | ggctacggtctccGTAAAACCGTTACCTTTGGAGAGACTACAGTGAAAGAAATACCGCCACCTGATGTTGTGCCTATTAAGGTTAGCATAGAGTGTTGTGGTGAACCATGGAACACTATCTTCAAGAAAGCTTATAAAGAGCCCATTGAAGTTGAAACTGATCTCACAGTAGAACAATTGCTCTCTGTGATCTATGAGAAAATGTGCGACGATCTCAAATTGTTTCCGGAGGCACCAGAACCTCCACCTTTTGAGAATGTCGCACTTGTTGATAAGAACGGGAAAGATTTGGATTGTATAAAATCATGCCACCTTATCTACCGTGATTGTGAGAGCGATGATGACATCGAGGAAGAAGATGCTGAAGAATGTGACACAGATTCAGCTGATGCTGAGGAGTGTGACACCGCCTCAGAATGTGAAGAAGAGGACGAGGACACTAAAGTGCTGGCTCTTCTACAGGACCCAGCAAGTAATAAGTACCCCTTACCTCTTGATGAAGACTATAGCGTTTACAATGGGTGCATTGTACATAAAGACGCTCTTGATGTTGTGAATCTACCATCTGGTGAAGAAACCTTCGTTGTCAACAACTGTTTTGATGGAGCAGTTAAACCGCTGCCACAAAAAGTTGTT |

| | |
|---|---|
| | GACGTTCTCGGCGATTGGGGTGAGGCTGTTGATGCACAAGAACAACTGTGTCAACAGGAACACGCTCAA<br>AACAATTCTGAAGGACCAGTGGAAAATTCTACTGATGATTCTAGTGTGTCGACTGAACAAGTTGTTGTTG<br>AAGAGAAGGAATTAATTCCCGTTGTTGAAGACGAACAAGCGGTAGTTGTTTACACCCCTGTTGAGCTACA<br>AGTTGTTGATGAAACACCAGATGAGTTTATTCTCACTGCTGATGTTCCTACAGAAGAATTTGTGTCTCAAG<br>AAGAAAAGGAGCCACAGGTCGAACAAGAGCATGTTCAGGTTGCTAAAGCGCAACGTGAAAGGAAGGC<br>TAAAAAGTTCAAGGTTAAACTAACTACATGTGAAAAACCCAAGTATTTGGAGTATAAAACATGTGTAGGTG<br>ATTTGACTGTAGTAGTTGCCAAAGCATTGGATGAGTTTAAAGAATTCTGCCTTGTAAACGCTGCAAATGAG<br>CATATGTCTCATGGGGGCGGTGTTGCAAAGGCAATTGCGGACTTTTGTGGACCAGATTTTGTAGAATATT<br>GTGCGGACTATGTTAAGAAACATGGGCCACAGCAACGACTTGTCACACCTTCATATGTTAGAGGCATTCA<br>ATGTGTGAACAATGTTGTTGGACCACGCCATGGAGACAGTAACTTGGAGGATAAACTCGTTGCTGCATAC<br>AAGAATGTGCTTGTAGACGGTGTTGTAAATTATGTTGTTCCTGTTCTCTCATCAGGAATCTTTGGTGTAGAC<br>TTTAAATTGTCAATAGACGCAATGCGCAAAGCCTTTGAGGGTTGTGACATACGCGTTTTGCTCTTCTCTCT<br>GAGTCAAGAACACATCGAGTACTTTGATGCAACTTGTAAACAGAAGACAATTTATCTTACAGATGATGGT<br>GTGAAATACCGCTCTGTTATTGTGAAACCTGGTGACTCTTTGAGTCAATTTGGACAGGTTTTTGCAAAGA<br>ATAAGACAGTTTTTACAGCAGATGATGTAGATGATAAAGAAATTCTCTTTACGCCTACTACTGACAAGACT<br>GTTCTTGATTATTACCATTTAGATGCGCAAAAGTATGTGATATATTTACAAACTCTTGCGCAGAAGTGGAAT<br>GTCCAGTATAGGGACAATTTTATTGTATTGGAGTGGCGTGATGGAAATTGCTGGGTTAGTTCAGCAATAGT<br>ACTCCTTCAAGCTGCTAAGGTGAAGTTTAGAGGTTATCTTGCAGAAGCATGGGCCAAGTTCTTAGGTGGA<br>GATCCTACAGAATTTGTTGCATGGTGTTATGCGAGTTGCAATGCTAAAGTTGGTGAATTTTCAGATGCCAA<br>TTGGCTTTTGGCTAATTTGGCAGAATATTTTGATGCAGATTACACGAACGCATTTCTTAAGAAGCGCGTAT<br>CATGTAACTGTGGTGTTAAGAGTTATGAGCTTAGGGGCCTTGAAGCTTGTATTCAACCAGTAAGGGCACC<br>TAACCTTCTACATTTTAAGACGCAATATTCAAATTGCCCAATCTGTGGCGCAAATAGTACGGATGAAGTAAC<br>AGAAGCTTCATTACCATACTTATTGCTCTTTGCTACTGATGGTCCAGCTACAGTTGATTGTGATGAAAATGC<br>TGTAGGGACTGTTGTTTTTATTGGATCCACTAATAGTGGCCATTGTTATACACAGGCCTTTGGTAAGGCTTT<br>TGATAATCTTGCTAATGATAGAAAATTTGGAAAGAAGTCGCCATACATTACAACAATGTATACGCGATTCTC<br>TCTCAagagaccgtagcc |
| D388-F4 | ggctacggtctctCTCAAGAGTGAAAATCCGCTGTCTGAAAAGAGTAAGGATAAAACTAAAGTAGTAAAAGA<br>AGATGTTTCTAACCTTGCCACTAGTTCTAAGGCCAGTTTTGATGATCTTACTGACTTTGAACGGTGGTATG<br>ATAGTAACATCTATGAAAGTCTTAAAGTGCAGGAAATACCTGAGAAATTGGATGGTTATGTGTCATTTACA<br>ATAAAGGAAGATTCTAAGTTGCCGTTGACACTTAAAGTTAGAGGTATTAAATCAGTTGTTGACTTTAAGTC<br>GAAGAACGGTTTCACTTATAAGTTAACACCTGACACTGATGAAAATTCAAAAGCACCAATTTACTATCCAG<br>TCTTGGACTCTATTAGTCTTAAGGCAATATGGGTGGAAGGTGGTGCTAATTTTGTTGTTGGTCATCCAAAT<br>TATTATAGTAAATCTCTCCGCATTCCTACTTTTTGGGAAAGTGCAGAGAGTTTTGTTAAAATGGGTGATAAA<br>GTTGCTGGTGTAACTATGGGCCTTTGGCGTGTAGAGCATCTTAACAAACCTAATCTGGAGAGAATTTTCAA<br>CATTGCTAAGAAAGCTATTGTTGGATCTAGTGTTGTTACTACGCAATGTGGTAAAGTAATTAGTAAGGCAG<br>CTACATTCATTGCTGATAAAATAGGTGGGGGTGTAGTTCGCAACATTGTAGATAGAATTAAGGGTATTTGT<br>GGGTTTACACGTGGGCATTTTGAAAGAAAATTGTCTCCACAATTCATAAAAACACTTATATTCTTTTTCTTT<br>TACTTTATAAAGGCTAGTGCTAAGGGTATAGTTGCTAGCTACAAAAGTGTGTTATGTAAGGTGGTATTTGCT<br>ACTTTATTTATAGTGTGGTTTGTATACACAAGTAATCCAGTAATGTTTACTGGAATACATGTGCTAGATTTCC<br>TATTTGAAGGTTCTTTGTGTGGTCCTTATAATGACTACGGTAAAGATTCTTTTGATGTGCTACGCTATTGTG<br>GAGATGATTTTACTTGTCGTGTGTGTTTACATGACAGAGACTCTCTACATTTGTATAAACATGCTTATAGCG<br>TAGAACAGATTTATAAAGTTGCAAGTTCTGGCCTTGGTTTTAATTGGAATTGGCTTTATTTGGTCTTTCTAA<br>TCTTATTTGTTAAGCCAGTGGCAGGTTTTGTTATTATTTGCTATTGTGTTAAGTTTTTAGTGTTGAGTTCAAC<br>TGTGCTCCAAACTGGTGTAGGTTTTCTAGACTGGTTTATTCAGACAGTTTTTACACACTTTAATTTTATGGG<br>AGCAGGATTTTATTTCTGGCTCTTTTATAAAGTATACATACAGGTGCATCATATACTGTATTGTAAGGATGTA<br>ACATGTGAAGTGTGCAAAAGAGTTGCACGCAGCAATAGACAAGAGGTTAGCGTGGTTGTTGGTGGACG<br>CAAGCAAATAGTGCATGTCTACACTAACTCTGGCTATAATTTTTGTAAGAGCCATAATTGGTATTGTAGAAA<br>TTGTGATGGTTATGGTCACCAAAATACATTTATGTCTCCGGAAGTTGCTGGCGAGCTTTCTGAAAAGCTTA<br>AGCGCCATGTTAAACCTACATCATATGCTTACCACGTTGTGGATGAAGCATGCGTAGTTGACGGCTTTGTT<br>AATTTAAAATATAAAGCTGCAATTCCTGGTAAGGATAGTGCATCTATTGCTGTTAAGTGTTTCAGTGTTACA<br>GATTTCTTGAAGAAAGCTGTTTTTCTTAAGGAAGCATTGAAATGTGAACAAATATCCAATGATGGTTTTAT<br>AGTGTGTAATACACAGAGTTCGCATGCATTGGAGGAAGCAAAGAATGCAGCCATCTATTATGCGCAATGT |

| | |
|---|---|
| | CTGTGTAAACCAATACTTATACTTGACCAGTTACTTTATGACCAATTAGTAGTAGAGCCTGTGTCTAAGAGT GTTATAGATAAAGTTTGTAGTATTTTGTCTAATATAATATCTGTAGATACTGCAGCTTTAAATTATAAAGCAG GCACACTTCGTGATGTTCTGCTTTCTATTACTGGAGATGAAGAAGCCGTAGATATGGCTATCTTCTGTCATA ATCATGATGTGGAGTATACTAGTGATGGTTTTACTAATGTGATACCATCATATGGTATAGACACTGATAAATT AACACCTCGTGATAGAGGGTTTTTAATAAATGCAGATGCTTCTATTGCTAATTTGAGAGTTAAAAATGCTCC ACAGGTAGTATGGAAGTTTTCAGATCTTATTAAGTTGTCTGACAGTTGCCTTAAATATCTAATTTCAGCTAC TGTCAAGTCAGGATGTCGTTTCTTTATAACAAAGTCTGGTGCTAAACAAGTTATTTCTTGTCATACCCAGAA ATTGTTGGTAGAGAAAAAGGCTGGTGGTGTTGTCAACAagagaccgtagcc |
| D388-F5 | ggctacggtctctAACAGCACTTTTAAATCGGTTAAGAGTTGTCTTAAATGGCTTTTTGTCTTCTATATACTCTTT ACAGCATGTTGTTTGGGTTATTACTATATGGAGATGAATAGAAGTTTTGTTCATCCCATGTATGATTTTAATG CTACACTACATGTTGAAGGGTTTAAGGTTATTGATAAAGGTGTTATTAGAGAAATTGTATCAGAGGATAAT TGTTTCTCTAATAAGTACACTAATTTTGATGCATTTTGGGGTAAACCATATGTTAATAGTAGAGAGTGTCCA ATTGTTACAGCTGTCATAGATGGCGCTGGAACAATAGCAGCTGATGTTCCTGGTCTTGTATACTGGGTTCT CGATGGTGCTATGTTTATACACATGGCACAAACAGAAAGAAAACCGTGGTATATTCCTACTTGGTTTAATA GAGAAATTGTTGGTTACACCCACGATTCTATTATTACAGAAGGTGAGTTTTATACTTCCATAGCATTGTTTG CTTCTAGATGTTTGTATTTAACATCTAGTAACACACCACAGCTTTATTGCTTTAATGGTGATAATGATGCTCC TGGTGCTCTACCATTTGCAAGTATACTTCCTCATAGAGTATACTTTCAACCTAATGGAGTCAGGCTTATTGT GCCTCAGCAAATTATGCACACACCATACGTAGTTAAGTTAGTTTCAGACAGCTATTGTAGAGGTAGTGTTT GTGAGGCTACTAAACCAGGTTATTGTGTCTCTATGAACTCTCAATGGGTTTTGTTTAATGATGAATACACAA TTAAACCAGGTGTATTTTGTGGTTCTACTGTTAGAGAACTTTTGTTTAGTATGGTTAGCACATTTTTTACGG GTGTTAATCCAAATATTTATATGCAACTGGCAACTATGTTCTTAATACTTGTTGCCGTTGTGTTAGTTTTTGC AATGGTTATAAAGTTTCAAGGTGTTTTTAAAGCTTATGCAACTATTGTATTTATTATAATGCTTGTTTGGTTT ATTAATGCATTTGTATTATGTGTACATAGTTATAATAGTGTTTTAGCTATTATACTACTTGTGCTCTATTGTTAT GCATCACTTGTGACGAGTCGTAATACTGCCATAATAATGCACTGTTGGCTCGTTTTTACATTTGGTTTAATA GTACCTAATTGGATAGCTTGTGTCTACTTAGGTTTTGTGCTATACATGTACACACCGCTGTTTTTGTGGTGC TATGGTACAACTAAAAATTGCCGCAAGCTTTATGAAGGCAATGATTTTGTTGGTAATTATGATCTTGCTGCG AAGAGTACGTTTGTAATTCGCGGCCCTGAATTTGTTAAACTTACTAATGAGATAGGTGATAAATTTGAACA TTATCTCTCAGCGTATGCTAGGCTTAAGTACTACTCTGGCACTGGCAGTGAGCAAGATTACTTGCAAGCCT GTCGTGCATGGTTGGCCTATGCACTAGACCAATTTAGAAGCAGTGGTGTGGAAGTAGTTTATACTCCACCA CGATATTCAATTGGTGTTAGTAGATTACAAGCTGGTTTTAAGAAACTAGTATTTCCTAGTAGTGCTGTTGAG AAGTGCATTGTTAGTGTATCCTATAGGGGTAATAATCTTAATGGATTGTGGTTGGGTGACTCCATCTACTGT CCACGTCATGTGTTAGGTAAGTTTAGTGGTGACCAATGGGGTGATGTACTTAATCTTGCTAATAATCATGA GTTTGAAGTTGTAACTGGAAATGGTGTTACTTTGAGTGTTGTCAGTAGGCGTTTGAAAGGTGCAGTTTTA ATTTTACAAACTGCAATTGTTAATGCTGAAACTCCAAAGTACAAATTTATGAAAGCAAATTGTGGTGATAG TTTTACTATTGCTTGCTCTTATGGAGGTACAGTTATAGGACTTTACCCCGTTACTATGCGTTCTAATGGAACT ATTAGAGCATCTTTTCTTGCTGGAGCATGCGGTTCAGTAGGTTTTAATATAGAAAAGGGTGTAGTAAATTT TTACTACATGCACCATCTAGAGTTACCTAATGCATTACATACAGGGACTGACCTAATGGGTGAGTTCTATGG TGGCTATATAGATGAAGAGGTTGCTCAGAAAGTTCAACCCGATAAATTAGTTACTAATAATATTTTAGCATG GCTTTATGCAGCAATTATTAGTGTTAAAGAGAGTAGTTTTTCAACACCTAAGTGGCTTGAAAGTACTACTG TCAGTATTGAAGATTATAATAAGTGGGCAGTGGATAATGGTTTTACATCATTTGTAAGCTGCACTGCTATTA CTAAATTAAGTGCTATAACAGGAGTTGATGTTTGTAAACTCCTTCGCACTATTATGGTAAAAAGTACACAAT GGGGTAGTGACCCTATTTTGGGACAATATAATTTTGAAGATGAAATGACACCAGAATCTGTTTTTAATCAG GTGGGTGGTGTTAGGtgagaccgtagcc |
| D388-F6 | ggctacggtctcaTAGGTTACAGTCTTCTGTTGTAAAGAAAGCTGCGTCTTGGTTTTGGAGTAGATGTGCGCT AGCTTGCTTTTTATTTGTCTTGTGTTCTATTGTGCTGTTCACAGCCTTACCATATAGATATTATTTATATGGCG CTGCTGTTTTATTTGCAGCTGTGCTCTTTATTTCATTTACTGTGAAACATGTTATGGCATTTATGGATACTTTC CTCTTGCCAACACTGATTACAGTTATTATTGGAGTTTGTGCTGAAGTACCTTTTATCTACAATACTCTAATTA GTCAAATTGTTATTTTCTTTAGCCAATGGTATGATCCAGTAGTCTTTGACACTGTAGTACCGTGGATGTTTCT ACCATTAGTCTTGTACACAGCTTTTAAATGTATACAAGGTTGTTATAGTGTAAACTCCTTTAATACTTCTCTG TTAGTGTTGTATCAGTTTATGAAGTTGGGTTTTGTTATATATACCTCTTCTAATACTCTTACAGCCTACTCAGA AGGTAATTGGGAGTTATTCTTTGAGTTAGTGCACACAACTGTGTTGGCTAATGTTAGTAGTAATTCTTTAAT TGGTTTAATTGTGTTTAAATTTGCTAAGTGGATGCTGTATTATTGTAATGCCTCATACCTTAATAATTATGTCC |

| | |
|---|---|
| | TAATGGCTGTCATTGTTAATGGCATAGGTTGGATGTTTACTTGTTACTTTGGATTTTATTGGTGGATTAATAA<br>GGTCTTTGGTTTAACCTTAGGTAAATATAGTTTCAAAGTTTCAGTAGATCAATATAGGTATATGTGTCTTCAT<br>AAGATTAACCCACCTAAAACTGTGTGGGAAGTCTTTTCTACAAATATACTTATACAAGGTATAGGTGGTGA<br>CCGTGTGTTGCCTGTAGCTACAGTGCAATCTAAATTGAGTGATGTAAAGTGTACAACTGTTGTCTTAATGC<br>AGCTTTTGACTAAGCTTAATGTTGAAGCAAACTCAAAAATGCATGTTTATCTTGTAGAGTTACACAATAAA<br>ATTCTTGCCTCAGATGATGTTAATGAGTGCATGGATAACTTATTGGGTATGCTTGTAACACTATTTTGTGTAG<br>ATAGTACTATTGATTTAAGTGAGTATTGTGATGATATACTTAAGAGGTCTACTGTTTTACAGTCAGTCACTCA<br>AGAGTTTTCACACATACCTTCTTATGCAGAGTATGAAAGAGCTAAAGACCTCTATGAAAAGGTTTTAGCTG<br>AGTCTAAAAATGGTGGTGTTACACAGCAAGAGCTTGCTGCATATCGTAAAGCTGCCAATATTGCAAAGTC<br>AATCTTTGATAGAGACTTGGCTGTTCATAAGAAGTTGGATAGCATGGCTGAACGTGCTATGACAACTATGT<br>ATAAGGAAGCGCGTGTTACTGATAGAAGAGCTAAACTTGTTTCGTCATTACATGCGCTACTATTTTCTATGC<br>TTAAGAAGATAGATTCTGAAAAGCTTAATGTACTATTTGACCAGGCAAGTAGTGGTGTTGTACCGCTAGCT<br>ACTGTTCCAATAGTTTGTAGTAATAAGCTTACTCTTGTAGTGCCAGATCCAGAGACGTGGATCAAGTGCGT<br>AGAGGGCATGCATGTTACATATTCAACAGTTGTTTGGAATATTGATAATGTCATTGATGCAGATGGCACTG<br>AATTGCAACCAATTTCTACAGGTAATGGTTTAATTTACTGTATAAGTGGTGATAATATAGCATGGCCTCTTAA<br>GGTCAATTTGACTAGGAATGTGCATAATAAAGTTGATGCAGTATTGCAGAACAACGAGCTTATGCCTCATG<br>GTGTTAAAACAAAGGCATGCGTGGCAGGTGTAGATCAAGCACACTGCAGCGTAGAGTCTAAATGTTATTA<br>TACTAATATTAGTGGTAATTCAGTAGTGGCTGCTATTACCTCTTTAAATCCTAATTTGAAAGTTGCCTCATTT<br>TTAAATGAAGCAGGCAATCAAATTTATGTTGACTTAGACCCACCATGTAAATTTGGCATGAAGGTGGGTG<br>ATAAAGTAGAAGTTGTTTACTTGTATTTTATAAAGAATACAAGGTCAATTGTCAGAGGTATGGTACTTGGT<br>GCTATATCTAATGTTGTGGTTTTACAGTCTAAAGGATATGAGACAGAGGAAGTTGATGCTGTTGGCATACT<br>ATCACTTTGTTCTTTTGCAGTAGATCCGGCTGACACGTATATTAAATATGTGGCTGCAGGTAACCAACCTTT<br>AGGTAACTGTGTAAAAATGTTGACAGTTCATAATGGTAGTGGCTTTGCTATAACATCAAAGCCAAGCCCAA<br>CTCCTGACCAGGATTCTTACGGAGGGGCTTCTGTGTGTCTCTATTGCAGGGCACATATAGCTCACCCGGG<br>AAGTGCAGGAAATTTAGATGGACGTTGTCAATTTAAAGGTTCTTTTGTACAAATACCTACTACGGAGAAA<br>GATCCTGTAGGGTTCTGTCTACGTAATAAGGTTTGCACTGTTTGTCAGTGTTGGATAGGTCATGGATGTCA<br>ATGTGATTCACTCAGACAACCAAAACCTTCAGTTCAATCAGATGCTGGcgagaccgtagcc |
| D388-F6-10mt<br>(nsp10.P85L) | ggctacggtctcaTAGGTTACAGTCTTCTGTTGTAAAGAAAGCTGCGTCTTGGTTTTGGAGTAGATGTGCGCT<br>AGCTTGCTTTTTATTTGTCTTGTGTTCTATTGTGCTGTTCACAGCCTTACCATATAGATATTATTTATATGGCG<br>CTGCTGTTTTATTTGCAGCTGTGCTCTTTATTTCATTTACTGTGAAACATGTTATGGCATTTATGGATACTTTC<br>CTCTTGCCAACACTGATTACAGTTATTATTGGAGTTTGTGCTGAAGTACCTTTTATCTACAATACTCTAATTA<br>GTCAAATTGTTATTTTCTTTAGCCAATGGTATGATCCAGTAGTCTTTGACACTGTAGTACCGTGGATGTTTCT<br>ACCATTAGTCTTGTACACAGCTTTTAAATGTATACAAGGTTGTTATAGTGTAAACTCCTTTAATACTTCTCTG<br>TTAGTGTTGTATCAGTTTATGAAGTTGGGTTTTGTTATATATACCTCTTCTAATACTCTTACAGCCTACTCAGA<br>AGGTAATTGGGAGTTATTCTTTGAGTTAGTGCACACAACTGTGTTGGCTAATGTTAGTAGTAATTCTTTAAT<br>TGGTTTAATTGTGTTTAAATTTGCTAAGTGGATGCTGTATTATTGTAATGCCTCATACCTTAATAATTATGTCC<br>TAATGGCTGTCATTGTTAATGGCATAGGTTGGATGTTTACTTGTTACTTTGGATTTTATTGGTGGATTAATAA<br>GGTCTTTGGTTTAACCTTAGGTAAATATAGTTTCAAAGTTTCAGTAGATCAATATAGGTATATGTGTCTTCAT<br>AAGATTAACCCACCTAAAACTGTGTGGGAAGTCTTTTCTACAAATATACTTATACAAGGTATAGGTGGTGA<br>CCGTGTGTTGCCTGTAGCTACAGTGCAATCTAAATTGAGTGATGTAAAGTGTACAACTGTTGTCTTAATGC<br>AGCTTTTGACTAAGCTTAATGTTGAAGCAAACTCAAAAATGCATGTTTATCTTGTAGAGTTACACAATAAA<br>ATTCTTGCCTCAGATGATGTTAATGAGTGCATGGATAACTTATTGGGTATGCTTGTAACACTATTTTGTGTAG<br>ATAGTACTATTGATTTAAGTGAGTATTGTGATGATATACTTAAGAGGTCTACTGTTTTACAGTCAGTCACTCA<br>AGAGTTTTCACACATACCTTCTTATGCAGAGTATGAAAGAGCTAAAGACCTCTATGAAAAGGTTTTAGCTG<br>AGTCTAAAAATGGTGGTGTTACACAGCAAGAGCTTGCTGCATATCGTAAAGCTGCCAATATTGCAAAGTC<br>AATCTTTGATAGAGACTTGGCTGTTCATAAGAAGTTGGATAGCATGGCTGAACGTGCTATGACAACTATGT<br>ATAAGGAAGCGCGTGTTACTGATAGAAGAGCTAAACTTGTTTCGTCATTACATGCGCTACTATTTTCTATGC<br>TTAAGAAGATAGATTCTGAAAAGCTTAATGTACTATTTGACCAGGCAAGTAGTGGTGTTGTACCGCTAGCT<br>ACTGTTCCAATAGTTTGTAGTAATAAGCTTACTCTTGTAGTGCCAGATCCAGAGACGTGGATCAAGTGCGT<br>AGAGGGCATGCATGTTACATATTCAACAGTTGTTTGGAATATTGATAATGTCATTGATGCAGATGGCACTG<br>AATTGCAACCAATTTCTACAGGTAATGGTTTAATTTACTGTATAAGTGGTGATAATATAGCATGGCCTCTTAA<br>GGTCAATTTGACTAGGAATGTGCATAATAAAGTTGATGCAGTATTGCAGAACAACGAGCTTATGCCTCATG |

| | |
|---|---|
| | GTGTTAAAACAAAGGCATGCGTGGCAGGTGTAGATCAAGCACACTGCAGCGTAGAGTCTAAATGTTATTA TACTAATATTAGTGGTAATTCAGTAGTGGCTGCTATTACCTCTTTAAATCCTAATTTGAAAGTTGCCTCATTT TTAAATGAAGCAGGCAATCAAATTTATGTTGACTTAGACCCACCATGTAAATTTGGCATGAAGGTGGGTG ATAAAGTAGAAGTTGTTTACTTGTATTTTATAAAGAATACAAGGTCAATTGTCAGAGGTATGGTACTTGGT GCTATATCTAATGTTGTGGTTTTACAGTCTAAAGGATATGAGACAGAGGAAGTTGATGCTGTTGGCATACT ATCACTTTGTTCTTTTGCAGTAGATCCGGCTGACACGTATATTAAATATGTGGCTGCAGGTAACCAACCTTT AGGTAACTGTGTAAAAATGTTGACAGTTCATAATGGTAGTGGCTTTGCTATAACATCAAAGCCAAGCCCAA CTCCTGACCAGGATTCTTACGGAGGGGCTTCTGTGTGTCTCTATTGCAGGGCACATATAGCTCACCTGGG AAGTGCAGGAAATTTAGATGGACGTTGTCAATTTAAAGGTTCTTTTGTACAAATACCTACTACGGAGAAA GATCCTGTAGGGTTCTGTCTACGTAATAAGGTTTGCACTGTTTGTCAGTGTTGGATAGGTCATGGATGTCA ATGTGATTCACTCAGACAACCAAAACCTTCAGTTCAATCAGATGCTGGcgagaccgtagcc |
| D388-F7 | ggctacggtctcgCTGGTGCACCTGGTTTTGATAAGAATTATTTAAACGGGTACGGGGTAGCAGTGAGGCTCG GCTGATACCCCTTGCTAATGGTTGTGAACCTGATGTTGTAGAGCGAGCCTTTGATGTATGTAATAAGGAAT CAGCCGGTATGTTTAAAAATTTGAAGCGTAACTGTGCTCGATTCCAAGAAGTGTGTGGTACTGAAGATGG AAATCTTGAGTATCGTGATTCTTATTTTGTGGTTAAACAAACCACTCCTAGTAATTATGAACATGAGAAAGC CTGTTATGGGGACTTAAAGTCAGAAGTAACAGCTGATCATGATTTCTTTGTGTTCAATAAGAACATTTATAA TATTAGTAGGCAGAGGCTTACTAAATATACTATGATGGACTTTTGTTATGCTTTGCGGCATTTTGACCCAAA GGATTGCGAAGTTCTTAAAGAAATACTTGTCACTTATGGTTGTATAGAAGATTACCATCCTAAGTGGTTCG AAGAAAATAAGGATTGGTACGATCCAATAGAAAACCCAAATTATGCCATGTTAATGGCTAAAATGGGACCT ATTGTTCGTCGTGCATTGTTGAATGCTGTTGAGTTTGGGAACCTAATGGTTGAGAAGGGTTATGTAGGTG TAGTTACACTAGATAACCAGGACCTCAATGGCAAATTTTATGATTTTGGTGATTTTCAGAAAACAGCACCT GGTGCTGGTGTTCCTGTTTTTGATACATATTATTCCTACATGATGCCCATCATAGCCATGACGGATGCGTTAG CACCTGAAAGGTATTTTGAATATGATGTGCATAAGGGTTACAAGTCTTATGACCTCCTCAAGTATGATTATA CAGAGGAGAAGCAAGAACTGTTTCAAAAGTACTTCAAGTACTGGGATCAAGAATACCACCCTAATTGTCG AGATTGTGTCGACGATAGGTGTTTGATACATTGTGCAAACTTCAACGTACTTTTCTCTACACTAATACCACA AACTTCTTTTGGTAATTTATGTAGAAAAGTGTTCGTTGATGGTGTACCATTTATAGCTACTTGTGGCTATCAC TCTAAGGAACTTGGTGTTATTATGAATCAAGATAACACCATGTCTTTTTCAAAAATGGGTTTAAGCCAGCTT ATGAAATTTGTAGGAGATCCCGCTCTTTTAGTAGGAACTTCTAACAATTTAGTTGATCTTAGAACTTCTTGC TTTAGTGTCTGTGCGTTAGCTTCTGGTATAATACACCAAACAGTTAAGCCAGGTCACTTTAATAAAGATTTC TACGATTTTGCTGAAAAAGCAGGTATGTTTAAGGAAGGATCATCAATTCCACTTAAACACTTCTTCTATCCA CAGACTGGTAATGCTGCTATAAACGATTATGATTATTATCGTTATAACAGGCCTACTATGTTTGATATACGCC AGCTTCTATTTTGTTTAGAGGTGACGTCTAAATATTTTGAATGTTATGAAGGTGGTTGTATACCAGCTAGTC AAGTGGTTGTTACCAACCTTGACAAGAGTGCAGGCTTTCCATTTAATAAATTTGGAAAAGCCCGTCTCTAT TATGAAATGAGTCTAGAAGAACAGGACCAACTCTTTGAAAGTACAAAGAAAAATGTCTTGCCTACTATAA CTCAGATGAATCTTAAATATGCTATATCCGCGAAAAATAGAGCGCGTACTGTGGCAGGTGTTTCCATTCTTT CTACTATGACTAACAGACAGTTCCATCAGAAGGTTCTTAAGTCTATAGTCAATACTAGAAATGCACCTGTAG TTATTGGAACAACTAAATTTTATGGCGGTTGGGACAATATGCTAAGAAACCTAGTTCAAGGTGTGGAAGA CCCAATTCTTATGGGTTGGGACTATCCTAAGTGTGATAGAGCAATGCCAAATTTGTTGCGTATAGCAGCGT CTTTGGTGCTTGCCCGTAAACATACTAATTGTTGCACATGGTCTGAGCGCATTTATAGATTGTACAATGAAT GCGCCCAAGTATTGTCTGAAACTGTATTAGCTACAGGTGGTATATATGTGAAACCTGGTGGTACTAGCAGT GGTGATGCCACCACTGCTTATGCCAACAGTGTTTTTAATATAATACAAGCCACATCTGCTAATGTTGCGCGT TTACTTAGTGTTATAACGCGTGATATAGTATATGATGACATTAAAAGCCTGCAGTATGAGTTATACCAGCAG GTCTATAGGCGAGTCAACTTTGATCCTGCCTTTGTTGAAAAGTTTTATTCTTACATGTGTAAGAATTTTTCA TTAATGATCTTGTCTGATGATGGTGTTGTTTGTTATAACAACACACTTGCCAGACAAGGTCTTGTAGCAGA CATTTCTGGTTTTAGAGAAATTCTCTACTACCAAAACAATGTTTATATGGCCGATTCTAAGTGTTGGGTTGA ACCCGACTTAGAAAAAGGCCCACATGAATTTTGTTCACAGCACACAATGCTAGTAGAGGTTGATGGCGA GCCAAGATACTTGCCATATCCAGACCCATCACGCATTTTGGGTGCATGTGTATTTGTAGATGAAGTGGATA AGACAGAACCTGTGGCTGTTATGGAGCGTTATATAGCTCTCGCCATAGATGCTTACCCTCTTGTACATCATG AAAATGAGGAGTACAAGAAAGTTTTCTTTGTACTCCTTTCTTATATTAGAAAGCTCTATCAGGAGCTCTCTC AAAGTATGCTTATTGACTATTCCTTTGTTATGGATATAGACAAGGGTAGTAAATTTTGGGAACAGGAGTTCT ATGAAAATATGTATAGGGCTCCTACAACTTTACAATCTTGTGGTGtgagaccgtagcc |

| | |
|---|---|
| D388-F8 | ggctacggtctcaGGTGTCTGTGTAGTTTGCAATAGTCAAACTATACTACGCTGTGGTAATTGTATTAGAAAACC<br>ATTCTTGTGTTGTAAGTGTTGCTATGACCATGTCATGCACACTGACCACAAAAATGTTTTGTCTATAAATCC<br>ATACATTTGCTCTCAACCTGGTTGTGGTGAAGCGGATGTTACTAAACTTTACCTTGGAGGTATGTCTTACTT<br>CTGTGGTAATCATAAACCTAAATTATCAATACCGTTAGTATCTAATGGTACTGTTTTTGGAATTTATAGGGCC<br>AACTGTGTAGGTAGCGAAAATGTTGATGATTTTAATCAACTAGCTACTACTAACTGGTCTACTGTTGAACCT<br>TATATATTAGCAAATCGTTGTAGCGACTCATTAAGACGCTTTGCTGCTGAGACAGTTAAAGCAACAGAAGA<br>GTTGCATAAACAGCAATTTGCTAGTGCAGAGGTCAGAGAGGTTATCTCTGACCGTGAATTAATTCTATCAT<br>GGGAACCTGGTAAGACTAGGCCTCCCTTGAATAGAAATTATGTGTTTACTGGCTATCACTTTACCAGGACA<br>AGTAAGGTACAGCTAGGTGATTTTACTTTTGAGAAAGGTGAAGGTAGAGATGTTGTCTATTATAGGGCGA<br>CGTCTACAGCTAAATTGTCTCCTGGAGACATTTTTTTCTTAACATCACACAATGTTGTTTCTCTTGTAGCAC<br>CAACTTTGTGTCCTCAACAAACTTTTTCTAGGTTTGTTAATTTAAGACCTAATGTAATGGTACCGGAGTGTT<br>TTGTTAATAACATTCCATTATACCATTTAGTAGGTAAACAAAAGCGTACTACAGTACAAGGCCCTCCTGGCA<br>GTGGTAAGTCACACTTTGCCATAGGCCTTGCTGCATACTTTAGTAATGCTCGTGTGGTTTTTACAGCATGTT<br>CTCATGCTGCAGTAGATGCGCTTTGTGAAAAAGCGTTTAAATTTCTTAAAGTTGATGATTGCACTCGCATA<br>GTACCTCAAAGGACTACTGTTGACTGCTTCTCAAAATTTAAAGCTAATGACACAGGCAAAAAGTACATTTT<br>TAGTACTATTAATGCCTTGCCAGAAGTTAGTTGTGACATTCTGTTGGTAGATGAGGTTAGTATGTTGACCAA<br>TTATGAATTGTCATTTATTAATGGTAAGATAAATTACCAGTATGTTGTGTATGTAGGTGATCCTGCTCAATTAC<br>CGGCACCTCGTACTTTGCTTAATGGTTCACTTTCACCCAAGGATTATAATGTCATCACAAATCTTATGGTTT<br>GTGTTAAACCTGACATTTTCCTTGCGAAGTGTTACCGTTGTCCTAAAGAAATTGTAGACACTGTGTCTACT<br>CTTGTTTATGATGGAAAGTTTGTTGCAAACAACCCAGAATCACGTGAGTGCTTCAAGGTTGTAGTTAATAA<br>TGGTAATTCTGATGTAGGTCACGAGAGTGGTTCAGCCTACAACACAACACAATTGGAATTTGTGAAAAAT<br>TTTGTTTGTCGCAATAAACAATGGCGAGAAGCAACATTCATTTCACCTTATAATGCAATGAACCAGAGAGC<br>CTATCGTATGCTTGGACTTAATGTCCAGACGGTAGACTCCTCACAAGGTTCAGAGTATGATTATGTTATATT<br>CTGTGTTACAGCTGATTCTCCGCATGCGCTGAATATTAACAGATTCAATGTTGCGCTTACAAGAGCTAAGC<br>GTGGTATACTTGTTGTCATGCGCCAGCGTGACGAATTGTATTCAGCTCTTAAATTTACAGAGCTAGATTCTG<br>AGGCAATCCTGCAAGGTACAGGTTTGTTTAAAATTTGCAATAAGGAATTTAGTGGAGTACACCCTGCTTAT<br>GCAGTAACCACTAAAGCACTTGCGGCAACCTATAAGGTTAACGACGAGCTAGCAGCACTTGTTAATGTTG<br>AGGCTGGTTCAGAGATTACTTATAAACATCTTATTTCTCTGTTAGGGTTTAAGATGAGTGTTAATGTTGAAG<br>GTTGTCACAACATGTTTATAACACGTGATGAGGCAATTCGCAATGTAAGAGGATGGGTTGGTTTTGATGTA<br>GAAGCCACACATGCTTGTGGCACTAATATTGGCACTAACCTTCCATTTCAAGTAGGATTCTCCACTGGTGC<br>TGACTTTGTAGTCACACCTGAGGGTCTTATTGATACATCAATAGGCAATAATTTTGAGCCTGTGAACTCTAA<br>AGCACCTCCAGGTGAACAATTTAATCACTTGAGAGCTTTGTTTAAGAGTGCTAAGCCTTGGCATGTTATAA<br>GACCTAGGATAGTTCAGATGTTAGCAGACAATCTATGCAATGTTTCAGATTGTGTAGTGTTTGTCACATGG<br>TGTCATGGCCTAGAACTAACTACTTTGCGCTATTTTGTTAAAATAGGCAAGGAACAAGTTTGTTCTTGTGG<br>TTCTAGAGCTACAACTTTTAATTCTCATACTCAGGCTTACGCTTGTTGGAGGCACTGCCTGGGTTTTGATTT<br>TGTCTATAATCCACTTTTAGTGGATATTCAACAATGGGGTTACTCTGGTAATCTACAGTTTAATCATGATTTG<br>CATTGTAATGTGCATGGCCACGCTCATGTAGCTTCAGCGGATGCTATTATGACTCGCTGCCTTGCAATTAAT<br>AATGCATTTTGTAAAGATGTCAACTGGGAACTGCAGTATCCTCATATTGCAAATGAGGATGAAGTAAATTC<br>CAGTTGCAGGTATCTACAACGCATGTATCTTAATGCGTGTGTTGATGCTCTTAAAGTCAACGTTGTCTATGA<br>TATAGGCAACCCTAAAGGTATAAAATGTGTTAGACGTGGGGATGTCAATTTTAGGTTCTATGATAAGAACC<br>CAATTGTACCTAACGTCAAACAGTTTGAGTATGACTATAGTCAGCATAAAGATAAGTTTGCTGATGGTCTTT<br>GTATGTTTTGGAATTGTAATGTGGATTGTTATCCTGAAAATTCGTTAGTTTGTAGGTATGACACACGAAATT<br>TGAGTGTGTTTAATTTACCTGGTTGTAATGGTGGTAGCCTGTATGTTAATAAACATGCATTCCACACACCTA<br>AGTTTGATCGCATTAGCTTTCGTAATTTGAAAGCTATGCCATTCTTCTTCTATGACTCATCGCCCTGCGAAA<br>CTATTCAAGTTGATGGAGTAGCGCAAGACCTTGTGTCATTAGCAACTAAAGATTGTATCACAAAATGCAAC<br>ATTGGTGGTGCTGTTTGTAAGAAACATGCTCAAATGTATGCAGAATTTGTGGCTTCTTATAATGCAGCTGTT<br>ACAGCTGGTTTTACTTTTTGGGTTACTAATAATTTTAACCCTTACAACTTGTGGAAGAATTTTTCAGCACTC<br>CAagagaccgtagcc |
| D388-F8-14mt<br>(nsp14.V393L) | ggctacggtctcaGGTGTCTGTGTAGTTTGCAATAGTCAAACTATACTACGCTGTGGTAATTGTATTAGAAAACC<br>ATTCTTGTGTTGTAAGTGTTGCTATGACCATGTCATGCACACTGACCACAAAAATGTTTTGTCTATAAATCC<br>ATACATTTGCTCTCAACCTGGTTGTGGTGAAGCGGATGTTACTAAACTTTACCTTGGAGGTATGTCTTACTT<br>CTGTGGTAATCATAAACCTAAATTATCAATACCGTTAGTATCTAATGGTACTGTTTTTGGAATTTATAGGGCC |

| | |
|---|---|
| | AACTGTGTAGGTAGCGAAAATGTTGATGATTTTAATCAACTAGCTACTACTAACTGGTCTACTGTTGAACCT<br>TATATATTAGCAAATCGTTGTAGCGACTCATTAAGACGCTTTGCTGCTGAGACAGTTAAAGCAACAGAAGA<br>GTTGCATAAACAGCAATTTGCTAGTGCAGAGGTCAGAGAGGTTATCTCTGACCGTGAATTAATTCTATCAT<br>GGGAACCTGGTAAGACTAGGCCTCCCTTGAATAGAAATTATGTGTTTACTGGCTATCACTTTACCAGGACA<br>AGTAAGGTACAGCTAGGTGATTTTACTTTTGAGAAAGGTGAAGGTAGAGATGTTGTCTATTATAGGGCGA<br>CGTCTACAGCTAAATTGTCTCCTGGAGACATTTTTTTCTTAACATCACACAATGTTGTTTCTCTTGTAGCAC<br>CAACTTTGTGTCCTCAACAAACTTTTTCTAGGTTTGTTAATTTAAGACCTAATGTAATGGTACCGGAGTGTT<br>TTGTTAATAACATTCCATTATACCATTTAGTAGGTAAACAAAAGCGTACTACAGTACAAGGCCCTCCTGGCA<br>GTGGTAAGTCACACTTTGCCATAGGCCTTGCTGCATACTTTAGTAATGCTCGTGTGGTTTTTACAGCATGTT<br>CTCATGCTGCAGTAGATGCGCTTTGTGAAAAAGCGTTTAAATTTCTTAAAGTTGATGATTGCACTCGCATA<br>GTACCTCAAAGGACTACTGTTGACTGCTTCTCAAAATTTAAAGCTAATGACACAGGCAAAAAGTACATTTT<br>TAGTACTATTAATGCCTTGCCAGAAGTTAGTTGTGACATTCTGTTGGTAGATGAGGTTAGTATGTTGACCAA<br>TTATGAATTGTCATTTATTAATGGTAAGATAAATTACCAGTATGTTGTGTATGTAGGTGATCCTGCTCAATTAC<br>CGGCACCTCGTACTTTGCTTAATGGTTCACTTTCACCCAAGGATTATAATGTCATCACAAATCTTATGGTTT<br>GTGTTAAACCTGACATTTTCCTTGCGAAGTGTTACCGTTGTCCTAAAGAAATTGTAGACACTGTGTCTACT<br>CTTGTTTATGATGGAAAGTTTGTTGCAAACAACCCAGAATCACGTGAGTGCTTCAAGGTTGTAGTTAATAA<br>TGGTAATTCTGATGTAGGTCACGAGAGTGGTTCAGCCTACAACACAACACAATTGGAATTTGTGAAAAAT<br>TTTGTTTGTCGCAATAAACAATGGCGAGAAGCAACATTCATTTCACCTTATAATGCAATGAACCAGAGAGC<br>CTATCGTATGCTTGGACTTAATGTCCAGACGGTAGACTCCTCACAAGGTTCAGAGTATGATTATGTTATATT<br>CTGTGTTACAGCTGATTCTCCGCATGCGCTGAATATTAACAGATTCAATGTTGCGCTTACAAGAGCTAAGC<br>GTGGTATACTTGTTGTCATGCGCCAGCGTGACGAATTGTATTCAGCTCTTAAATTTACAGAGCTAGATTCTG<br>AGGCAATCCTGCAAGGTACAGGTTTGTTTAAAATTTGCAATAAGGAATTTAGTGGAGTACACCCTGCTTAT<br>GCAGTAACCACTAAAGCACTTGCGGCAACCTATAAGGTTAACGACGAGCTAGCAGCACTTGTTAATGTTG<br>AGGCTGGTTCAGAGATTACTTATAAACATCTTATTTCTCTGTTAGGGTTTAAGATGAGTGTTAATGTTGAAG<br>GTTGTCACAACATGTTTATAACACGTGATGAGGCAATTCGCAATGTAAGAGGATGGGTTGGTTTTGATGTA<br>GAAGCCACACATGCTTGTGGCACTAATATTGGCACTAACCTTCCATTTCAAGTAGGATTCTCCACTGGTGC<br>TGACTTTGTAGTCACACCTGAGGGTCTTATTGATACATCAATAGGCAATAATTTTGAGCCTGTGAACTCTAA<br>AGCACCTCCAGGTGAACAATTTAATCACTTGAGAGCTTTGTTTAAGAGTGCTAAGCCTTGGCATGTTATAA<br>GACCTAGGATAGTTCAGATGTTAGCAGACAATCTATGCAATGTTTCAGATTGTGTAGTGTTTGTCACATGG<br>TGTCATGGCCTAGAACTAACTACTTTGCGCTATTTTGTTAAAATAGGCAAGGAACAAGTTTGTTCTTGTGG<br>TTCTAGAGCTACAACTTTTAATTCTCATACTCAGGCTTACGCTTGTTGGAGGCACTGCCTGGGTTTTGATTT<br>TGTCTATAATCCACTTTTAGTGGATATTCAACAATGGGGTTACTCTGGTAATCTACAGTTTAATCATGATTTG<br>CATTGTAATGTGCATGGCCACGCTCATGTAGCTTCAGCGGATGCTATTATGACTCGCTGCCTTGCAATTAAT<br>AATGCATTTTGTAAAGATGTCAACTGGGAACTGCAGTATCCTCATATTGCAAATGAGGATGAAGTAAATTC<br>CAGTTGCAGGTATCTACAACGCATGTATCTTAATGCGTGTGTTGATGCTCTTAAAGTCAACGTTGTCTATGA<br>TATAGGCAACCCTAAAGGTATAAAATGTGTTAGACGTGGGGATGTCAATTTTAGGTTCTATGATAAGAACC<br>CAATTGTACCTAACGTCAAACAGTTTGAGTATGACTATAGTCAGCATAAAGATAAGTTTGCTGATGGTCTTT<br>GTATGTTTTGGAATTGTAATGTGGATTGTTATCCTGAAAATTCGTTACTTTGTAGGTATGACACACGAAATT<br>TGAGTGTGTTTAATTTACCTGGTTGTAATGGTGGTAGCCTGTATGTTAATAAACATGCATTCCACACACCTA<br>AGTTTGATCGCATTAGCTTTCGTAATTTGAAAGCTATGCCATTCTTCTTCTATGACTCATCGCCCTGCGAAA<br>CTATTCAAGTTGATGGAGTAGCGCAAGACCTTGTGTCATTAGCAACTAAAGATTGTATCACAAAATGCAAC<br>ATTGGTGGTGCTGTTTGTAAGAAACATGCTCAAATGTATGCAGAATTTGTGGCTTCTTATAATGCAGCTGTT<br>ACAGCTGGTTTTACTTTTTGGGTTACTAATAATTTTAACCCTTACAACTTGTGGAAGAATTTTTCAGCACTC<br>CAagagaccgtagcc |
| D388-F9 | ggctacggtctctTCCAGTCTATTGACAATATTGCTTATAATATGTATAAGGGTGGGCATTACGATGCTATTGCAG<br>GCGAAATACCCACAGTCATAACTGGAGATAAAGTTTTTGTTATTGATCAAGGTATTGAAAAGGCAGTTTTT<br>GTTAATCAAACAACTCTGCCTACATCAGTTGCATTTGAGTTGTATGCGAAGAGAAACATTCGCACACTGCC<br>AAATAACAGAATATTGAGTGGTTTAGGTGTAGACGTTACCCATGGTTTTGTAATCTGGGATTACGCTAACC<br>AAACGCCACTGTATCGTAATACTGTTAAAGTATGTGCATATACAGACATTGAACCTAATGGTTTAATAGTTCT<br>GTATGATGATAGATGTGGTGATTATCAATCTTTTCTTGCTGCTGATAATGCTGTTTTAGTGTCTACACAGTGT<br>TATAAGCGGTATCCTTATGTAGAAATACCATCAGATCTGCTTGTTCAGAATGGTATGCCACTAAAAGATGGA<br>GCGAATCTGTATGTTTATAAGCGGAGTAATGGAGCGTTTGTAACGCTACCAACCACATTAAACACACAGG |

| | |
|---|---|
| | GTCGTAATTATGAAACTTTTGAACCTCGTAGTGATGTTGAGAGAGATTTTCTTGACATGTTGGAGGACGAT<br>TTTATTGAAAAGTATGGTAAGGACCTAGGTCTACAACACATACTTTATGGTGAAGTAGAAAAACCACAATT<br>GGGTGGTTTACACACTGTTATAGGAATGTACAGACTCTTGCGTGCTAATAAATTAGACGCGAAGTCTGTAA<br>CCAATTCAGATTCTGATGTCATGCAAAACTATTTTGTTTTGGCAGACAATGGTTCTTACAAACAGGTTTGC<br>ACAGTTGTAGACCTTTTGCTTGATGACTTCTTAGAACTCCTTAGGAACATACTTAAAGAGTATGGTACTAAT<br>AAGTCCAAAGTTGTAACAGTGTCTATTGATTACCATAGCATAAATTTTATGACTTGGTTTGAAGAAGGCAG<br>TATTAAAACATGTTATCCACAGCTTCAGTCGGCTTGGACATGTGGTTATAATATGCCTGAACTTTATAAAGTT<br>CAGAACTGTGTTATGGAACCTTGTAATATTCCAAACTATGGTGTTGGAATTACATTGCCAAGTGGTATTATG<br>ATGAATGTGGCAAAGTATACACAACTTTGCCAATACCTCTCTAAAACCACAATGTGTGTACCGCATAACATG<br>CGAGTAATGCATTTTGGAGCTGGAAGTGATAAAGGAGTTGCCCCTGGCAGCACTGTCCTTAAACAGTGG<br>CTTCCAGAAGGAACACTCCTTGTTGATAACGATATTGTAGATTATGTATCTGATGCACACGTGTCTGTGCTG<br>TCAGACTGCAATAACTATAAAACAGAGCACAAGTTTGATCTTGTGATATCTGATATGTATACAGATAATGATT<br>CAAAAAGAAAACATGAAGGCATAGTAGCCAATGATGGCAATGACGATGTCTTCATTTATCTAGCTAATTTT<br>TTAAAGAACAACTTAGCTCTGGGTGGTAGTTTTGCCATCAAATTAACAGAGACAAGTTGGCATGAGAGTC<br>TTTATGATATTGCACAGGATTGTGCATGGTGGACAATGTTTTGTACAGCTGTGAATGCTTCATCTTCAGAA<br>GCATTCTTGCTCGGTATTAATTATTTGGGTGACAGTGGAAAAGTGAAAGTTAGTGGAAAAACACTGCACG<br>CAAATTATATATTTTGGAGGAATTGTAATTATTTACAAACTTCTGCTTATAGTGCATTTGATGTTGCTAAGTT<br>TGGATTGAAATTAAAAGCAACACCAGTTGTTAATTTGAAGAAGGAACAAAAGACCGACTTAGTAGTTAAT<br>TTACTAAGGAACGGTAAATTATTGGTTAGAGATtgagaccgtagcc |
| D388-F10 | ggctacggtctcaAGATGTTGGTGAAGTCACTGTTTTTAGTGACCATTTTGTGTGCACTATGTAGTGCAAATTT<br>GTTTGATTCTGATAATAATTATGTGTACTACTACCAAAGTGCTTTTAGACCGCCAAATGGGTGGCACCTACA<br>AGGAGGTGCTTATGCAGTAGTCAATTCTACTAATTATACTAATAATGCCGGTTCTGCACATGTGTGCACTGT<br>TGGTGTTATTAAGGATGTTTATAATCAAAGTGTGGCTTCCATAGCTATGACAGCACCTCTTCAGGGTATGGC<br>TTGGTCTAAGTCACAATTCTGTAGTGCACACTGTAACTTTTCTGAAATTACAGTTTTTGTCACACATTGTTAT<br>AGTAGTGGTAGTGGGTCTTGTCCTATAACAGGCATGATTCCACGTGATCATATTCGTATTTCTGCAATGAAA<br>AATGGTTCTTTATTTTATAATTTAACAGTTAGCGTATCTAAATACCCTAATTTTAAATCTTTTCAATGTGTTAA<br>CAACTTCACATCTGTTTATTTAAATGGTGATCTTGTTTTTACTTCCAACAAAACTACTGATGTTACGTCAGCA<br>GGTGTGTATTTTAAAGCAGGTGGACCTGTAAATTATAGTATTATGAAAGAATTTAAGGTTCTTGCTTACTTT<br>GTTAATGGTACAGCACAAGATGTAATTTTGTGCGACAATTCCCCCAAGGGTTTGCTAGCTTGTCAATATAA<br>CACTGGCAATTTTTCAGATGGCTTTTATCCTTTTACTAATAGTACTTTGGTTAGGGAAAAGTTCATCGTCTA<br>TCGCGAAAGTAGTGTTAATACTACTCTGGCGTTAACTAATTTCACTTTTACTAATGTAAGTAATGCACAGCC<br>TAATAGTGGTGGTGTTAATACTTTTCATTTATACCAAACACAAACAGCTCAGAGTGGTTATTATAATTTTAAT<br>TTGTCATTTCTGAGTCAGTTTGTGTATAAGGCAAGTGATTTTATGTATGGGTCCTACCACCCTAGTTGTTCT<br>TTTAGACCAGAAACCATTAATAGTGGTTTATGGTTTAATTCCTTGTCAGTTTCTCTTACCTATGGACCCCTAC<br>AGGGAGGGTGTAAGCAATCTGTTTTTAGTGGTAAGGCAACGTGTTGTTACGCTTACTCTTATAAAGGCCC<br>AATGGCATGTAAAGGTGTTTATTCAGGTGAATTAAGCACGAATTTTGAATGTGGATTGCTGGTTTATGTTA<br>CTAAGAGTGATGGCTCTCGTATACAGACTAGAACAGAGCCCTTAGTATTAACGCAATACAATTATAATAATA<br>TTACTTTAGATAAGTGTGTTGCCTATAATATATATGGCAGAGTAGGCCAAGGTTTTATTACTAATGTGACTGA<br>TTCTGCTGCTAATTTTAGTTATTTAGCAGATGGTGGGTTAGCTATTTTAGATACGTCGGGTGCCATAGATGT<br>TTTTGTTGTACAGGGCATCTATGGTCTTAATTATTACAAGGTTAATCCTTGTGAAGATGTTAATCAACAATTT<br>GTAGTGTCTGGTGGCAATATAGTTGGCATTCTTACTTCTAGAAATGAAACAGGTTCTGAACAGGTTGAGA<br>ACCAGTTTTATGTTAAGTTAACCAATAGCTCACATCGTCGTAGGCGTTCTATTGGCCAAAATGTAACAAGTT<br>GTCCTTATGTTAGTTATGGCAGATTTTGTATTGAACCAGATGGTTCGTTAAAGATGATAGTGCCAGAAGAA<br>TTGAAACAGTTTGTGGCACCTTTACTTAATATTACTGAAAGTGTACTCATACCTAACAGTTTTAATCTTACTG<br>TTACAGATGAGTACATACAAACACGTATGGATAAGGTCCAAATCAATTGCCTTCAATATGTTTGCGGCAATT<br>CTTTGGAGTGTAGAAAATTGTTTCAACAATATGGTCCGGTTTGTGATAATATATTGTCTGTTGTAAATAGTG<br>TTAGTCAAAAAGAAGATATGGAACTTTTAAGCTTCTATTCTTCTACTAAACCAAAGGGTTATGATACACCA<br>GTTCTTAGTAATGTAAGCACTGGTGAATTTAATATTTCTCTTCTCTTGAAACCCCCAAGTAGTCCTAGTGGG<br>CGTTCTTTCATTGAAGATCTTTTATTTACAAGTGTTGAAACAGTTGGTTTGCCAACTGATGCTGAATATAAA<br>AAATGCACAGCGGGACCTTTGGGTACTCTTAAAGATCTTATCTGTGCTAGGGAATATAATGGTTTATTAGT<br>GTTGCCTCCAATTATTACGGCGGATATGCAAACAATGTATACTGCTTCTTTAGTGGGTGCTATGGCCTTTGG<br>TGGTATTACATCAGCTGCAGCTATACCTTTTGCTACTCAGATTCAGGCAAGAATTAATCATCTTGGTATTACA |

| | |
|---|---|
| | CAGTCTTTGTTAATGAAAAATCAAGAAAAGATTGCTGCTTCCTTTAATAAGGCCATTGGTCATATGCAGGA<br>AGGTTTTAGAAGCACTTCGCTAGCATTACAACAGATTCAAGATGTTGTTAATAAGCAGAGTGCTATTCTTA<br>CTGAAACTATGAATTCTCTTAATAAGAATTTTGGTGCTATTACATCAGTCATTCAAGATATTTACGCGCAACT<br>TGATGCAATTCAAGCAGATGCACAAGTTGACCGCCTTATTACTGGTAGACTTTCATCACTCTCAGTGCTAG<br>CCTCTGCTAAACAGTCTGAGTATATTAGAGTTTCCCAGCAGCGTGAATTAGCCACTCAAAAAATTAATGAG<br>TGTGTTAAATCACAATCTAATAGGTACGGATTTTGTGGTAGTGGAAGACATGTTCTTTCGATACCACAAAAT<br>GCACCTAATGGTATAGTGTTTATACACTTTACTTATACACCAGAGAGTTTTGTTAATGTTACTGCAATAGTGG<br>GTTTTTGTGTAAATCCTGCTAATGCTAGTCAGTATGCTATAGTACCTGCTAATGGAAGGGGTATTTTTATACA<br>AGTTAATGGCACGTACTATATCACTGCACGTGATATGTATATGCCACGAGACATTACTGCAGGAGATATAGT<br>TACTCTTACGTCTTGTCAAGCAAATTATGTTAATGTAAATAAAACCGTCATTACTACATTTGTAGAAGATGAC<br>GATTTTGATTTTGATGATGAGTTGTCAAAATGGTGGAATGACACTAAGCATCAGCTACCAGACTTTGACGA<br>CTTCAATTACACAGTACCCATACTTAATATTAGCGGTGAAATTGATTATATTCAAGGTGTTATACAGGGTCTT<br>AATGACTCCCTTATAAACCTTGAAGAACTTTCAATAATTAAAACTTATATTAAGTGGCCTTGGTATGTTTGG<br>CTTGCCATAGGCTTTGCTATTATTATTTTTATCCTTATTTTAGGATGGGTGTTTTTCATGACTGGTTGTTGTG<br>GTTGTTGTTGTGGATGCTTTGGTATCATTCCTCTAATGAGTAAGTGTGGTAAGAAATCCTCTTATTACACGA<br>CTTTTGATAATGATGTGGTAACTTAACAATcgagaccgtagcc |
| D388-F10-Beau(S) | ggctacggtctcaAGATGTTGGTGAAGTCACTGTTTTTAGTGACCATTTTGTGTGCACTATGTAGTGCTGTTTT<br>GTATGACAGTAGTTCTTACGTTTACTACTACCAAAGTGCCTTCAGACCACCTAGTGGTTGGCATTTACAAG<br>GGGGTGCTTATGCGGTAGTTAACATTTCTAGCGAATTTAATAATGCAGGCTCTTCATCAGGGTGTACTGTT<br>GGTATTATTCATGGTGGTCGTGTTGTTAATGCTTCTTCTATAGCTATGACGGCACCGTCATCAGGTATGGCT<br>TGGTCTAGCAGTCAGTTTTGTACTGCACACTGTAATTTTTCAGATACTACAGTGTTTGTTACACATTGTTATA<br>AACATGGTGGGTGTCCTTTAACTGGCATGCTTCAACAGAATCTTATACGTGTTTCTGCTATGAAAAATGGC<br>CAGCTTTTCTATAATTTAACAGTTAGTGTAGCTAAGTACCCTACTTTTAGATCATTTCAGTGTGTTAATAATTT<br>AACATCCGTATATTTAAATGGTGATCTTGTTTACACCTCTAATGAGACTATAGATGTTACATCTGCAGGTGTT<br>TATTTTAAAGCTGGTGGACCTATAACTTATAAAGTTATGAGAGAAGTTAAAGCCCTGGCTTATTTTGTTAAT<br>GGTACTGCACAAGATGTTATTTTGTGTGATGGATCACCTAGAGGCTTGTTAGCATGCCAGTATAATACTGG<br>CAATTTTTCAGATGGCTTTTATCCTTTTACTAATAGTAGTTTAGTTAAGCAGAAGTTTATTGTCTATCGTGAA<br>AATAGTGTTAATACTACTTGTACGTTACACAATTTCATTTTTCATAATGAGACTGGCGCCAACCCTAATCCTA<br>GTGGTGTTCAGAATATTCAAACTTACCAAACAAAAACAGCTCAGAGTGGTTATTATAATTTTAATTTTTCCT<br>TTCTGAGTAGTTTTGTTTATAAGGAGTCTAATTTTATGTATGGATCTTATCACCCAAGTTGTAAATTTAGACT<br>AGAAACTATTAATAATGGCTTGTGGTTTAATTCACTTTCAGTTTCAATTGCTTACGGTCCTCTTCAAGGTGG<br>TTGCAAGCAATCTGTCTTTAAAGGTAGAGCAACTTGTTGTTATGCTTATTCATATGGAGGTCCTTCGCTGTG<br>TAAAGGTGTTTATTCAGGTGAGTTAGATCATAATTTTGAATGTGGACTGTTAGTTTATGTTACTAAGAGCG<br>GTGGCTCTCGTATACAAACAGCCACTGAACCGCCAGTTATAACTCAAAACAATTATAATAATATTACTTTAA<br>ATACTTGTGTTGATTATAATATATATGGCAGAACTGGCCAAGGTTTTATTACTAATGTGACCGACTCAGCTGT<br>TAGTTATAATTATCTAGCAGACGCAGGTTTGGCTATTTTAGATACATCTGGTTCCATAGACATCTTTGTTGTA<br>CAAGGTGAATATGGTCTTAATTATTATAAGGTTAACCCTTGCGAAGATGTCAACCAGCAGTTTGTAGTTTCT<br>GGTGGTAAATTAGTAGGTATTCTTACTTCACGTAATGAGACTGGTTCTCAGCTTCTTGAGAACCAGTTTTA<br>CATCAAAATCACTAATGGAACACGTCGTTTTAGACGTTCTATTACTGAAAATGTTGCAAATTGCCCTTATGT<br>TAGTTATGGTAAGTTTTGTATAAAACCTGATGGCTCAATTGCCACAATAGTACCAAAACAATTGGAACAGT<br>TTGTGGCACCTTTATTTAATGTTACTGAAAATGTGCTCATACCTAACAGTTTCAACTTAACTGTTACAGATG<br>AGTACATACAAACGCGTATGGATAAGGTCCAAATTAATTGCCTGCAGTATGTTTGTGGCAGTTCTCTGGAT<br>TGTAGAAAGTTGTTTCAACAATATGGGCCTGTTTGCGACAACATATTGTCTGTAGTAAATAGTGTTGGTCA<br>AAAAGAAGATATGGAACTTTTGAATTTCTATTCTTCTACTAAACCGGCTGGTTTTAATACACCAGTTCTTAG<br>TAATGTTAGCACTGGTGAGTTTAATATTTCTCTTCTGTTAACAAATCCTAGTAGTCGTAGAAAGCGTTCTCT<br>TATTGAAGACCTTCTATTTACAAGCGTTGAATCTGTTGGACTACCAACAAATGACGCATATAAAAATTGCAC<br>TGCAGGACCTTTAGGCTTTTTTAAGGACCTTGCGTGTGCTCGTGAATATAATGGTTTGCTTGTGTTGCCTC<br>CTATCATAACAGCAGAAATGCAAGCTTTGTATACTAGTTCTCTAGTAGCTTCTATGGCTTTTGGTGGTATTAC<br>TGCAGCTGGTGCTATACCTTTTGCCACACAACTGCAGGCTAGAATTAATCACTTGGGTATTACCCAGTCAC<br>TTTTGTTGAAGAATCAAGAAAAAAATTGCTGCTTCCTTTAATAAGGCCATTGGTCATATGCAGGAAGGTTTT<br>AGAAGTACATCTCTAGCATTACAACAAATTCAAGATGTTGTTAGTAAACAGAGTGCTATTCTTACTGAGAC<br>TATGGCATCACTTAATAAAAATTTTGGTGCTATTTCTTCTGTGATTCAAGAAATCTACCAGCAATTTGACGC |

| | |
|---|---|
| | CATACAAGCAAATGCTCAAGTGGATCGTCTTATAACTGGTAGATTGTCATCACTTTCTGTTTTAGCATCTGC<br>TAAGCAGGCGGAGTATATTAGAGTGTCACAACAGCGTGAGTTAGCTACTCAGAAAATTAATGAGTGTGTT<br>AAGTCACAGTCTATTAGGTACTCCTTTTGTGGTAATGGACGACATGTTCTAACCATACCGCAAAATGCACCT<br>AATGGTATAGTGTTTATACACTTTTCTTATACTCCAGATAGTTTTGTTAATGTTACTGCAATAGTGGGTTTTT<br>GTGTAAAGCCAGCTAATGCTAGTCAGTATGCAATAGTGCCCGCTAATGGTAGGGGTATTTTTATACAAGTT<br>AATGGTAGTTACTACATCACTGCACGAGATATGTATATGCCAAGAGCTATTACTGCAGGAGATGTAGTTACG<br>CTTACTTCTTGTCAAGCAAATTATGTAAGTGTAAATAAGACCGTCATTACTACATTCGTAGACAATGATGATT<br>TTGATTTTAATGACGAATTGTCAAAATGGTGGAATGATACTAAGCATGAGCTACCAGACTTTGACAAATTC<br>AATTACACAGTACCTATACTTGACATTGATAGTGAAATTGATCGTATTCAAGGCGTTATACAGGGTCTTAAT<br>GACTCTCTAATAGACCTTGAAAAACTTTCAATACTCAAAACTTATATTAAGTGGCCTTGGTATGTTTGGCTT<br>GCCATAGGCTTTGCTATTATTATTTTTATCCTTATTTTAGGATGGGTGTTTTTCATGACTGGTTGTTGTGGTT<br>GTTGTTGTGGATGCTTTGGTATCATTCCTCTAATGAGTAAGTGTGGTAAGAAATCCTCTTATTACACGACTT<br>TTGATAATGATGTGGTAACTTAACAATcgagaccgtagcc |
| D388-F11 | ggctacggtctcgCAATACAGACCTAAAAAGTCTGTTTAATGATACAGAAGCCAACATCTTTTCTAATAGTGTTA<br>ATTCTTCTTTGGTTTAAACTTGTGCTAAGTTGTTTTAGAGAGTGTGTGTTAGCACTCCTACAACTAATACGA<br>GTTCTACTCCAAATTATTAATAGTAATTTACAATCTAGGATGCTCCTTTGGCACAGCCTAGACTAATGTTAGA<br>TTTTGAGAAAACAATTGAAACAGGTGAAGTAGTAGTACAACAAATCAGTTTCAATTTACAACATATTTCAA<br>GTGTTCTAGAAACACAAATTTTTGACCCATTTGAGTGTTGTTACTATTCAAGTGGTAGTTTTTATGAAATAG<br>ATTCAGCTGACGACTTTTCAGATGATGAGTTTACTGAATAAATCGCTAGAAGAGAACGGCAGTTTTCTAAC<br>AGCCCTTTATATATTTGTAGCATTTGTAGCACTTTATCTTTTAGGTAGAGCGCTTCAAGCATTTGTACAAGC<br>AGCTGATGCTTGTTGTTTATTTTGGTATACATGGGTAGTAGTTCCAGGAGCTAAGGGTACAGCCTTTGTGT<br>ATAAACATACATATGGTAGAAAACTTAACAATCCGGAATTGGAACAGGTTATTGTTAACGAGTTCCCAAAG<br>AACGGTTGGAATAATAAAAACCCTGCAATTTTTCAAGATGTCGAACGGCACGGAAAATTGCACTCTTAAT<br>ACTGAACAAGCAGTTCAGCTTTTTAAAGAGTATAATCTATTCATAACCGCATTCCTTTTGTTTCTAACCATAC<br>TACTTCAGTATGGATACGCAACTAGGAGTCGGTTTATTTACATACTGAAAATGGTAGTGTTATGGAGCTTCT<br>GGCCCCTTAACATTGCAGTAGGTGTAATTTCATGTGTATACCCACCAAACACGGGAGGTCTTGTCGCAGC<br>GATAATACTTACTGTGTTTGCATGTTTTTCTTTTGTAGGTTATTGGATCCAGAGTTTTAGACTCTTTAAGCGT<br>TGTAGGTCATGGTGGTCATTTAACCCAGAGTCTAATGCCGTAGGTTCAATACTCCTTACAAATGGTCAACA<br>ATGCAATTTTGCTATAGAGAGTGTGCCAATGGTGCTTTCTCCAATTATAAAGAATGGTGTTCTTTATTGTGA<br>GGGTCAGTGGCTTGCTAAATGTGAACCAGACCATTTGCCTAAAGACATTTTTGTGTGTACACCAGATAGA<br>CGCAACATTTATCGTATGGTGCAGAAATATACTGGTGACCAAAGCGGAAATAAGAAAAGGTTTGCTACATT<br>TGTCTATGCAAAGCAGTCAGTAGACACTGGCGAGCTAGAAAGTGTAGCAACAGGAGGAAGTAGTCTTTA<br>CACATAAATGTGTGTGTGTAGAGAATATTTAAAATTATTCTTTGACAGTGCCTCTGTTTTAAGAGCGCGGA<br>AGAGTATTATTTTTGAGGATATTAATATAAACCCTCTTTGTTTCATGCTCTCCTTTCAAGAGTTATTATTTAAA<br>AAACAGTTTTTCCACTCTTTTGTGCCAAAAACTGTTGTTGTTAACGGTGTAGCCTTCCAAGTAGATAATGG<br>AAAAGTTTACTATGAAGGAAAATCCATTTTTCAGAAAGGTTGTTGCAGGTTATGGTCCCAGTATAAGAGG<br>GATTAAAATAGTCAAACCACCAACAACAATTTCTTGTACTAAAGGCGTTTGGACTTACAAGCGCTTAAACT<br>AAACAAATACGGAATGAAATGGTTGTCTAGTTTAGGACGAGCATTCACTTCCTGTTATAAAGCCCTACTATT<br>AACTCAATTAAGAGTTTTAGATAGGTTAATTCTAGATCACGGACCGAGGCGTACTTTAAGTTGTGCCAGGC<br>GAGTGCTTTTAATTCAATTAGATTTAGTTTACAGGTTGGCTTATACGCCCACCCAATCGCTGGTATGAATAA<br>TAGTAAAGATAATCCTTTTCGCGGAGCAATAGCAAGAAAAGCGCGAATTTATCTGAGAGAAGGATTAGAT<br>TGTGTTTACTTTCTTAACAAAGCAGGACAAGCAGAGCCTTGTCCCGCGTGCACATCACTAGTATTCCAAG<br>GGAAAACTTGTGAGGGGCACATAGATAATAATAACTTGCTATCATGGCGAGCagagaccgtagcc |
| D388-F12 | ggctacggtctctGAGCGGTAAAGCAACTGGAAAGACAGACTCCCCCGCGCCAATCATCAAACTAGGAGGGC<br>CAAAACCACCTAAGGTAGGGTCATCTGGAAATGCATCTTGGTTTCAGTCCCTAAAGGCCAAGAAACTAAA<br>TTCACCTCCTCCTATGTTTGAAGGTAGTGGTGTTCCTGATAATGAAAATCTAAAATCTAGCCAGCAGCATG<br>GATACTGGAGACGCCAAGCCAGGTATAAGCAAGGCAAAGGCGGAAGAAAACCAGTCCCTGATGCATGG<br>TACTTCTATTATACAGGAACAGGACCAGCCGCTGACCTGAATTGGGGTGATTCTCAAGATGGTATAGTGTG<br>GGTTGCTGCCAAGGGTGCTGATGTAAAATCTAGATCCAATCAGGGTACAAGAGATCCTGATAAGTTTGAC<br>CAATTTCCACTACGGTTTTCAGATGGAGGACCTGATGGTAATTTCCGTTGGGATTTCATTCCTCTGAATCGT<br>GGTAGGAGTGGGAGGTCGACTGCAGCTTCATCAGCAGCATCTAGTAGAGTGCCATCTCGAGAAGGTTCA<br>CGTGGTCGTAGGAGTGGAGCTGAAGAAGATCTGATTGCTCGCGCGGCAAAGATTATTCAGGACCAGCAA |

| | |
|---|---|
| | AGGAAGGGTACGCGTATTACAAAGCAAAAGGCAGATGAGATGGCTCACCGTCGATTCTGTAAGCGTACC<br>ATTCCACCAGGTTATAGAGTAGATCAAGTATTTGGCCCTCGTACTAAAGGTAAGGAGGGAAATTTTGGTG<br>ATGACAAGATGAATGAGGAAGGTATTAAGGATGGGCGTGTTACAGCAATGCTCAACCTTACACCTAGCCC<br>ACATGCTTGTCTTTTTGGAAGTAGAGTGACGCCCAAGCTTCAACCAGATGGGCTTCACCTTAGATTTGAA<br>TTTACTACTGTGGTGCCTAGAGATGACCCGCAGTTTGATAATTATGTAAAGATTTGTGATGAGTGTGTTGAT<br>GGTGTAGGCACACGCCCAAAAGACGAAGTTGTAAGACCAAAATCACGCTCAAGTTCAAGACCTGCTACA<br>AGAGGAAATTCTCCAGCGCCAAAACAACAGCGCCAAAAGAAGGAGAAAAAGCCAAAGAAGCAGGATG<br>ATGAAGTGGATAAAGCATTGACCTCAGATGAGGAGAGGAACAATGCACAGCTGGAATTTGATGATGAAC<br>CCAAAGTGATTAATTGGGGTGATTCAGCACTTGGTGAAAATGAACTTTGAGTAACATAATGGACTTGCTA<br>CATTTCATGTGGCATTGCTGTTGTTATTACTTCTCCTGCTTTCTTACAATCATTATGCGTTTTGATTGTGATCA<br>CGTGCAGTATCTTAGTTGGTTTTCCTTGTTGATTGCAGTTTGTTTTCTTTCTGTGCTTATTGTTTTTATTCTG<br>CTTTATTCTTTGTTATATCGTAGAAGTTTATTAAAGGTTAAGGAAGATAGGCATGTAGCTTGATTACCTACAT<br>GTCTATCGCCAGGGAAATGTCTAATCTGTCTACTTAGTAGCCTGGAAACGAACGGTAGACCCTTAGATTTT<br>AATTTAGTTTAATTTTTAGTTTAGTTTAAGTTAGTTTAGAGTAGGTTTAAAGATGCCAGTGCCGAGGCCAC<br>GCGGAGTACGATCGAGGGTACAGCACTAGGACGCCCATTAGGGGAAGAGCTAAATTTTAGTTTAAGTTA<br>AGTTTAATTGGCTAATTATATTTAAAATTTATAGGCTAGTATAGAGTTAGAGCAAAAAAAAAAAAAAAAAA<br>AAAAAAAAAAAATTCCTGCGGCCGGGTCGGCATGGCATCTCCACCTCCTCGCGGTCCGACCTGGGCATC<br>CGAAGGAGGACGTCGTCCACTCGGATGGCTAAGGGAGAGCTCGGATCCGGCTGCTAACAAAGCCCGAA<br>AGGAAGCTGAGTTGGCTGCTGCCACCGCTGAGCAATAACTAGCATAACCCCTTGGGGCCTCTAAACGGG<br>TCTTGAGGGGTTTTTTGCTGAAAGGAGGAACTATATCCGGATCGAGATCCTCTAGAGTCGACGGGCCGCC<br>ATGATGACAATAAAGAATTAATTATTGTTCACTTTATTCGACTTTAACCCAGCGGCCGCGGCTACGGagaga<br>ccgtagcc |

Supplementary Figure 2. Predicted Assembly Fidelity

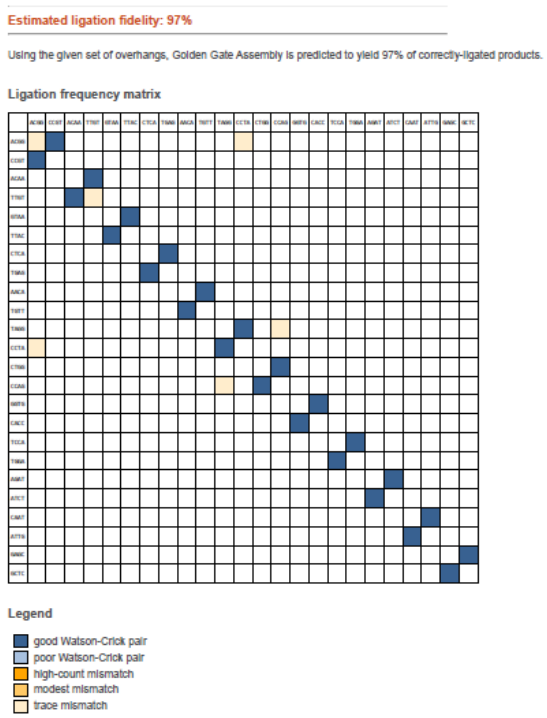


Data-optimized Assembly Design tools (ligasefidelity.neb.com) were used to check the fidelity of the assembly. The predicted ligation fidelity for the overhangs used in all Golden Gate Assemblies (ACGG, ACAA, GTAA, CTCA, AACA, TAGG, CTGG, GGTG, TCCA, AGAT, CAAT, GAGC) is 97%. The reaction conditions used for this prediction were “BsaI-HFv2 37-16 cycling.”

Supplementary Table 3. Component Fragments for each rIBV assembly.

| Fragment | D388-GGA | D388-14mt-GGA | D388-10.14mt-GGA | D388-Beau(S)-GGA |
|---|---|---|---|---|
| 1 | D388-F1 | D388-F1 | D388-F1 | D388-F1 |
| 2 | D388-F2 | D388-F2 | D388-F2 | D388-F2 |
| 3 | D388-F3 | D388-F3 | D388-F3 | D388-F3 |
| 4 | D388-F4 | D388-F4 | D388-F4 | D388-F4 |
| 5 | D388-F5 | D388-F5 | D388-F5 | D388-F5 |
| 6 | D388-F6 | D388-F6 | D388-F6-10mt | D388-F6 |
| 7 | D388-F7 | D388-F7 | D388-F7 | D388-F7 |
| 8 | D388-F8 | D388-F8-14mt | D388-F8-14mt | D388-F8 |
| 9 | D388-F9 | D388-F9 | D388-F9 | D388-F9 |
| 10 | D388-F10 | D388-F10 | D388-F10 | D388-F10-Beau(S) |
| 11 | D388-F11 | D388-F11 | D388-F11 | D388-F11 |
| 12 | D388-F12 | D388-F12 | D388-F12 | D388-F12 |

Supplementary Figure 3.

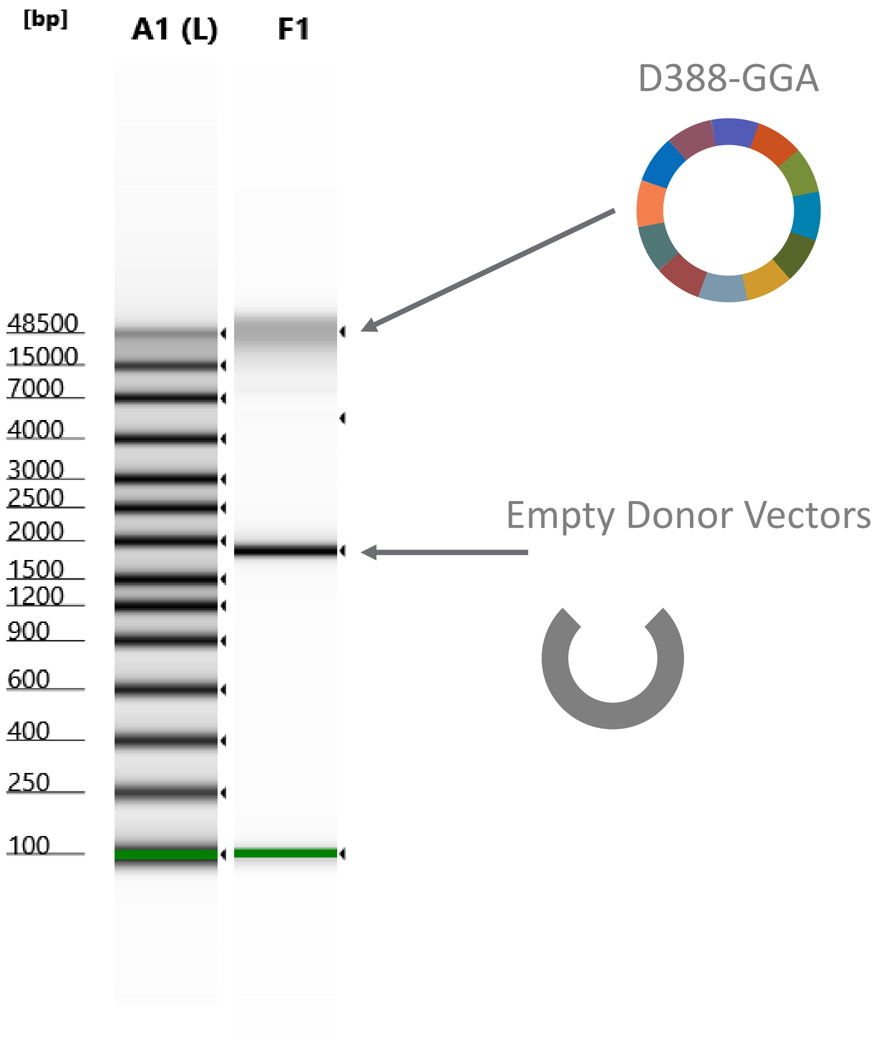


Assembly success was optionally assessed prior to rescue via visualization on Tapestation, a gel electrophoresis system that can visualize fragments up to ~40kb. The full length D388-GGA cDNA (28 kb) peak runs between the largest standards included in the kit, 15 kb and 48.5 kb. Under these cycling conditions, the majority of the DNA is seen in this full-length peak, with a second large peak at 1.9 kb that is the vector backbone cut away from the fragments during assembly.
